# Phosphorylation Deficient Inducible cAMP Early Repressor (ICER) Modulates Tumorigenesis and Survival in a Transgenic Zebrafish (*Danio rerio*) Model of Melanoma

**DOI:** 10.1101/2025.02.21.639575

**Authors:** Justin Wheelan, Melissa Spigelman, Angelo Cirinelli, James Reilly, Carlos A. Molina

## Abstract

Melanoma, the most lethal form of skin cancer, is commonly associated by mutations in the BRAF gene, particularly BRAF^V600E^, which drives tumor proliferation via ERK1/2 signaling cascade. While BRAF inhibitors initially demonstrate efficacy, therapeutic resistance remains a significant challenge. Emerging evidence implicates the cAMP signaling pathway, particularly the cAMP response element-binding protein (CREB) and its repressor, Inducible cAMP Early Repressor (ICER), in melanoma progression and drug resistance. ICER, a transcriptional repressor regulated via Ras/MAPK-mediated phosphorylation and ubiquitination, is degraded in melanoma, undermining its tumor-suppressive role. In a *braf^V600E^*; *p53*(loss of function) transgenic zebrafish (*Danio rerio*) model, we investigated the role of a ubiquitin-resistant ICER mutant (S35-41A-ICER) in tumor progression. Transgenic fish expressing S35-41A-ICER exhibited extended survival and reduced tumor invasiveness compared to wild-type ICER. RNA sequencing revealed dysregulation of CREB/CREM targets and compensatory pathways, including Rap1 and PI3K/AKT signaling, as well as candidate gene targets of ICER regulation, including the Protein Kinase A catalytic subunit *prkacaa*. Our findings suggest that a ubiquitin resistant ICER mitigates melanoma progression and represses oncogenic pathways in a *braf^V600E^* melanoma context.

**Summary Statement:** This study shows that ubiquitin-resistant ICER mutant suppresses melanoma progression, prolongs survival in *braf^V600E^* zebrafish, revealing its potential as a tumor suppressor and therapeutic target in melanoma resistance.

## Introduction

Melanoma, originating from pigment-producing melanocytes, remains the most lethal form of skin cancer despite its relative rarity compared to other cutaneous malignancies (Cancer.gov). In 2024 an estimated 100,640 new cases of invasive melanoma were diagnosed and 8,290 associated deaths were reported in the United States (Cancer.org, 2024).

Approximately 70–80% of melanocytic nevi and ∼50% of malignant melanomas harbor mutations in the *BRAF* gene, with BRAF^V600E^ being the most prevalent alteration. This mutation leads to constitutive activation of the extracellular signal-regulated kinase (ERK1/2) cascade, driving tumor proliferation and survival (Davies et al., 2002; Wan et al., 2004). Although pharmacological inhibition of BRAF^V600E^, using agents such as vemurafenib, has shown efficacy in reducing tumor burden, these responses are often transient due to the development of therapeutic resistance (Bollag et al., 2012; Chan et al., 2017). Immune checkpoint inhibitors, including anti-PD1 (nivolumab, pembrolizumab) and anti-CTLA-4 (ipilimumab), have revolutionized melanoma treatment, especially in late-stage metastatic disease, increasing long-term survival to 50% from less than 10% historically (Carlino et al., 2021). Despite these advances, treatment resistance continues to pose significant challenges, underscoring the urgent need for alternative therapies (Ascierto et al., 2024; Winder et al., 2017).

Emerging evidence highlights the importance of the cyclic AMP (cAMP) signaling pathway in melanoma progression and resistance, despite infrequent genetic mutations within this pathway (Johannessen et al., 2013; Gough 2013). Transcriptional regulation downstream of cAMP is mediated following phosphorylation of cAMP response element-binding protein (CREB) and other members of cAMP responsive proteins such as, cAMP Responsive Element Modulator (CREM), which includes activators and repressors that bind cAMP-response elements (CREs) promoter motifs via basic leucine zipper domains. One way the activation of these kinase inducible proteins occurs, is through cAMP induced activation of Protein Kinase A (PKA).

Secondary messenger activity via the cAMP-PKA signaling pathway has been implicated in numerous processes relevant to tumor biology, such as glycogen metabolism, ion channel regulation, cell differentiation and proliferation and gene induction (Rohlff & Glazer, 1995). Resistance to BRAF and MEK inhibitors in BRAF^V600E^ melanoma has been linked to compensatory activation of cAMP-dependent signaling networks, involving PKA, and CREB (Johannessen et al., 2013). Biopsies of BRAF^V600E^ melanoma suggest that phosphorylated (and therefore active) CREB is suppressed by RAF-MEK inhibition, but restored in relapsing tumors (Johannessen et al., 2013). We hypothesize that ICER (Inducible cAMP Early Repressor), an antagonist of CREB and CREM in this context, may play a critical role in melanoma resistance and tumor progression.

ICER arises from an intronic promoter of the *CREM* gene and exists as four isoforms: ICER-I, ICER-Iγ, ICER-II, and ICER-IIγ, all of which retain a DNA-binding domain but lack the kinase-inducible and N-terminal domains present in other CREB family members (Stehle et al., 1993; Molina et al., 1993). Despite its compact structure, ICER functions as a potent transcriptional repressor, acting as a homodimer or heterodimer with other CREB family members to autoregulate its own expression by binding the prototypical palindromic CRE sequences (5‘ TGACGTCA 3’) within its promoter (Molina et al., 1993). Evolutionarily conserved, ICER retains ∼85% sequence identity across species, underscoring its critical role as a nuclear effector of cAMP signaling.

ICER’s regulatory significance extends to diverse physiological processes, including circadian rhythm (Takahasi, 1994), cardiac myocyte regulation (Tomita et al., 2003), cell cycle regulation (Yehia et al., 2001), and as a tumor suppressor (Pigazzi et al., 2008; Mémin et al., 2002). However, its degradation and subcellular localization are tightly controlled via post-translational modifications. Our prior work demonstrated that ICER proteasomal degradation is influenced by Ras/MAPK-mediated phosphorylation and subsequent ubiquitination. Using Tyr/Tet-Ras *INK4a−/−* transgenic mice and melanoma-derived R545 cells, we showed that ICER degradation is dependent on oncogenic H-RasV12G expression (Memin et al., 2011).

Pharmacological inhibition of Ras or the proteasome abolished ICER degradation, highlighting the role of Ras-driven proteasomal regulation (Healey et al., 2013). Further studies revealed that ICER is contextually regulated by phosphorylation at serine’s 35 and 41. Phosphorylation at serine 35 leads to ICER monoubiquitination and cytosolic relocalization (Healey et al., 2013), while phosphorylation at serine 41 triggers polyubiquitination and proteasomal degradation (Memin et al., 2011). The latter process is mediated by the E3 ubiquitin ligase UBR4, operating via the N-end rule pathway and requiring recognition of ICER’s N-terminus (Cirinelli et al., 2022). Our prior studies revealed that a modified N-terminal ICER tagged with human influenza hemagglutinin (NHA-ICER) induced apoptosis five times more effectively than C-terminal HA- tagged ICER), despite no significant differences in DNA-binding affinity between the two variants (Cirinelli et al., 2022).

In this study, we aimed to investigate the impact of ICER phosphorylation and subsequent ubiquitination on melanoma progression using a *braf^V600E^* +/+, *p53* (loss-of-function), and *mitf* (loss-of-function) transgenic zebrafish (*Danio rerio*) model. In this model we employ the miniCoopR system, where *mitf* expression is restored only upon successful transgene integration. Using this model, we generated three distinct transgenic cohorts: (1) ubiquitin- resistant ICER with serine-to-alanine substitutions at positions 35 and 41, and an N-terminal hemagglutinin (HA) tag (S35-41A-ICER); (2) wild-type ICER with an N-terminal HA tag (wtICER); and (3) an enhanced GFP (EGFP) control. We evaluated overall survival of these cohorts over 104 weeks. Tumor histology was assessed via hematoxylin and eosin (H&E) staining. Additionally, tumor-derived cell lines were established to perform RNA sequencing and investigate ICER’s role in transcriptional regulation. Our findings demonstrate that fish expressing the ubiquitin-resistant S35-41A-ICER mutant exhibit extended survival compared to wtICER and EGFP controls. Histological analysis revealed that tumors expressing wtICER were the most aggressive and invasive, while tumors in the S35-41A-ICER cohort displayed markedly reduced invasiveness. Cell lines derived from wtICER tumors showed accelerated proliferation compared to EGFP controls, consistent with ICER’s inability to exert its repressive function due to cytosolic relocation. Interestingly, we were unable to establish stable cell lines from S35-41A- ICER tumors, likely due to its suppression of cell proliferation, reinforcing its role as a tumor suppressor. RNA sequencing revealed a set of differentially expressed genes between wtICER and EGFP cell lines, many of which overlap with known CREB/CREM target. Gene Ontology (GO) of differentially expressed genes suggests compensatory rewiring of cellular networks in response to ICER dysregulation leading to an increase in cell proliferation, focal adhesion, and pro-metastatic signaling pathways, such as Rap1 and PI3K/AKT. Finally, we also propose a novel mechanism, in which ICER may directly reduce cAMP activation of PKA and subsequent CREB phosphorylation via competitive CRE binding on the *prkacaa* promoter, encoding the Protein Kinase A catalytic α-subunit. In this article we highlight ICER’s dual role as a repressor of oncogenic pathways and a target of proteasomal degradation and show that the ubiquitin-resistant S35-41A-ICER mutant not only mitigates tumor progression but also prolongs survival in a *braf^V600E^* driven melanoma context.

## Results

### Expression of ICER protein is abnormal in zebrafish melanomas

To validate and extend previous findings from Healy et al. (2013), which demonstrated reduced ICER expression in human melanoma cells and a mouse model for melanomagenesis, we sought to confirm these observations in a zebrafish model. Endogenous ICER protein expression was assessed in tissue samples collected from the skin or melanomas of seven individual zebrafish. As shown in Figure 1, ICER protein expression was reduced in tissues containing melanomas (lanes 3–7). In contrast, ICER was strongly expressed in normal skin tissues from both wild-type AB fish and non-tumor transgenic fish (lanes 1 and 2). These results mirror the previously reported findings and suggest that ICER is abnormally degraded in tumor tissues. The melanomas in our zebrafish model arose from melanocyte-specific expression of *braf^V600E^*, a mutation known to constitutively activate the MAPK pathway, reenforcing the hypothesis that ICER may play a pivotal role in melanomagenesis and the maintenance of the transformed phenotype.

**Figure 1.**
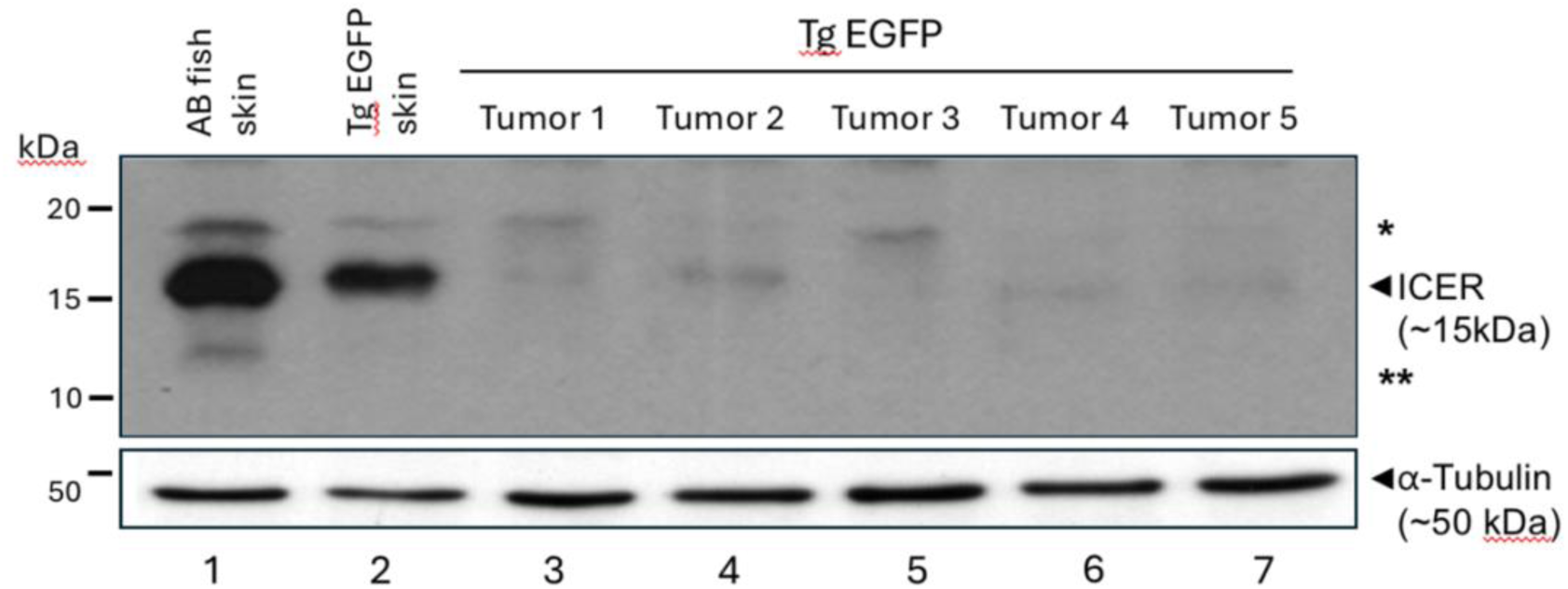
**ICER protein expression in zebrafish tissues**. Western blot analysis of protein expression in melanomas from the indicated zebrafish strains. Tissue samples were collected from skin tissues and tumors as shown. Western blot for endogenous ICER expression was performed from whole tissue extracts as described previously using anti-ICER polyclonal antibodies (Cirinelli et at. 2022). The same membrane was stripped and re-probed with anti ⍺- Tubulin antibodies as a loading control (bottom panel). The left margin indicates the relative mobility of Low Range Molecular Weight Markers from BioRAD (in kD). Indicated in the right margin is the relative mobility of ICER and ⍺-Tubulin. * and ** indicates post-translationally modified form of ICER (Memin et al. 2011; Cirinelli et at. 2022).

### ICER expression impacts tumorigenesis and survival in *braf* V600E, *p53* loss-of-function, and *mitf* loss-of-function transgenic zebrafish

We then tested whether autochthonous ICER expression in melanocytes would impact the survival of zebrafish with a constitutively active *braf^V600E^* mutation, in the context of a loss of function of *p53* gene, and with *mitf* rescued via the Tol2 integration of the miniCoopR plasmid, as described in Ceol et al., 2011 and Iyengar et al., 2012. Pair-wise log rank testing suggests a significant (p<0.05) *decrease* in survival in the fish, expressing wtICER (n=48) with an N-terminal HA-Tag, and demonstrable lower median and overall survival compared to EGFP control (n=37) (Fig 2.). Simultaneously, we evaluated the impact of the phosphorylation deficient ICER mutant (S35-41A-ICER) (n=40), also containing an N-terminal HA-tag.

**Figure 2.**
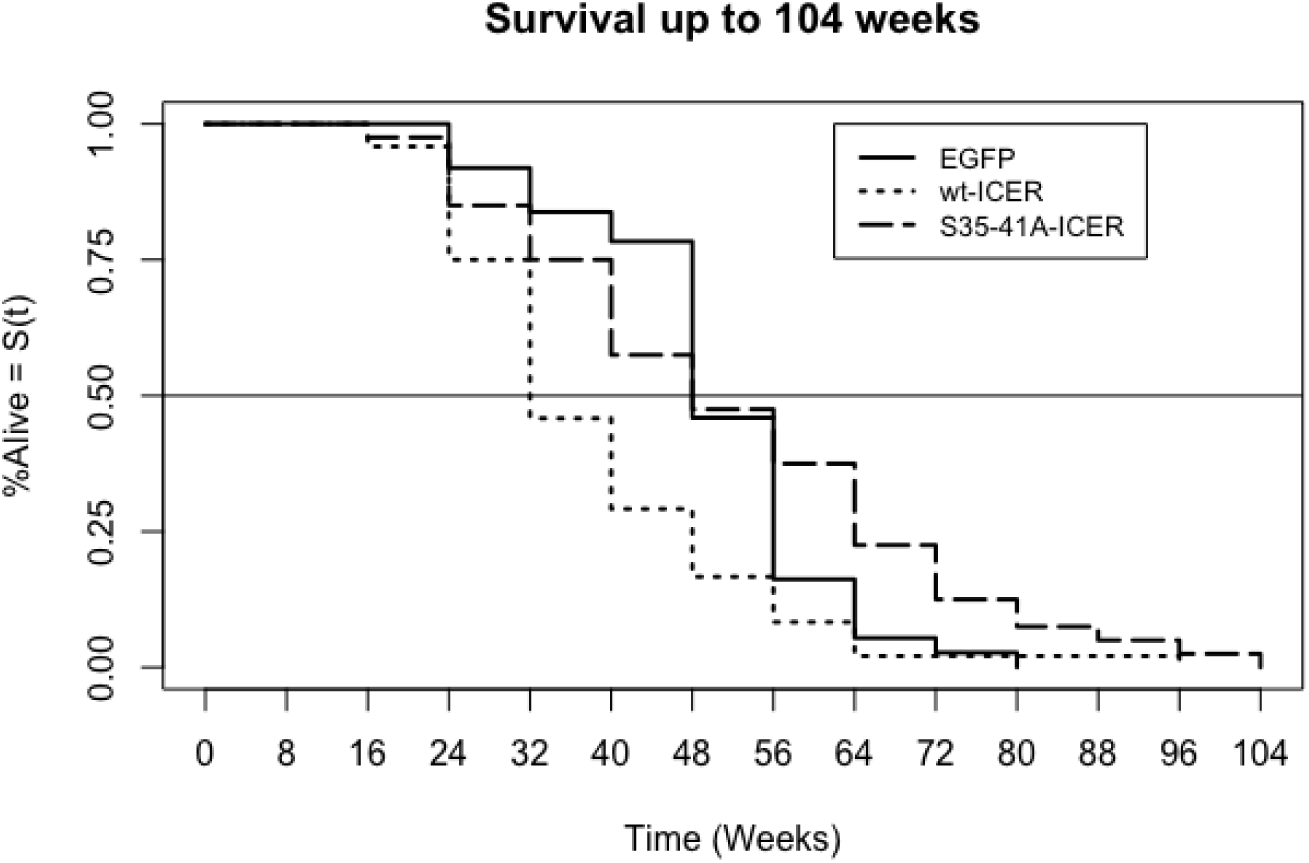
Survival Analysis. Kaplan-Meier survival curve comparing percent survival among three transgenic zebrafish cohorts: EGFP (n = 37), wtICER (n = 48), and S35-41A-ICER (n = 40) over a 104-week observation period. Survival curves were generated using the Kaplan-Meier estimator, with survival differences assessed using the log-rank test (p<0.05). Fish expressing wtICER displayed significantly reduced survival compared to the EGFP control, correlating with more aggressive tumor histology observed in Figure 3. In contrast, fish expressing the S35-41A-ICER mutant showed improved survival compared to wtICER (p <0.05), consistent with reduced tumorigenesis.

**Figure 3.**
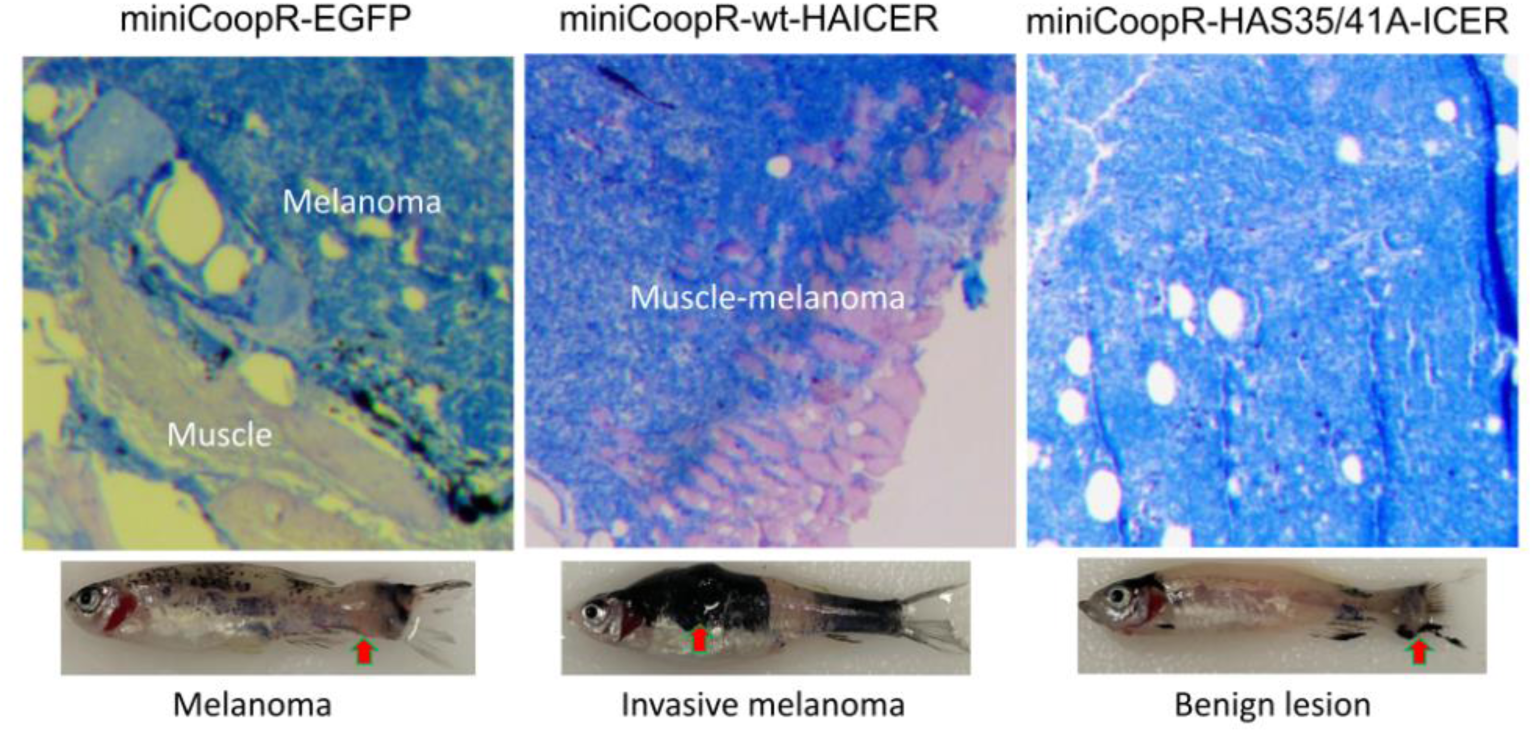
Tumor histology and gross phenotypic analysis of transgenic zebrafish cohorts expressing EGFP, wtICER, and S35-41A-ICER. Representative images of zebrafish and corresponding hematoxylin and eosin (H&E)-stained tumor sections for each cohort. From Left: EGFP-expressing zebrafish exhibit melanomas localized to the skin and minimal infiltration into proximal muscle tissue. Middle: wtICER- expressing zebrafish display extensive tumor growth with significant infiltration into the underlying muscle, indicative of an aggressive melanoma phenotype. Right: S35-41A-ICER expressing zebrafish show little to no tumor burden, with no evidence of melanoma invasion into the muscle, suggesting reduced malignancy. Gross phenotypic analysis mirrors these histological findings, with fish expressing EGFP showing moderate tumor size, wtICER fish exhibiting large, and widespread tumors, and S35-41A-ICER fish displaying minimal or potentially benign neoplastic tissue.

Conversely, this isoform rescued fish survival compared to the wtICER (p<0.05). However, while the S35-41A-ICER has a notably longer lifespan compared to both wtICER and EGFP control cohorts, the difference between S35-41A-ICER and EGFP control in this study was not statistically significant (p >0.05).

Histological evaluation of tumor progression and invasiveness was performed using hematoxylin and eosin (H&E) staining to visualize cellular and tissue architecture. The pathological analysis of the tumors shows that melanomas from EGFP-expressing fish show exophytic growth without invading the underlying musculature as shown before (Ceol et al., 2011). In contrast, the melanomas from wtICER-expressing fish invaded from the skin, through the collagen-rich stratum compactum of the dermis, into the underlying musculature. Fish expressing S35-41A-ICER do not show malignancies, showing signs of benign hyperplasia only, further supporting the hypothesis that the ubiquitin-resistant ICER variant mitigates tumor progression and invasion.

### Generation and Characterization of Zebrafish Tumor-Derived Cell Lines

To address the challenge of tumor heterogeneity in bulk RNA-seq experiments and to enable a more controlled analysis of transcriptional changes, we generated stable cell lines from tumors excised from transgenic zebrafish cohorts. Tumors were surgically extracted, enzymatically dissociated into single-cell suspensions, and cultured under conditions optimized for zebrafish cells (see Methods). These cell lines were derived from MiniCoopR-EGFP (control) and MiniCoopR-HA-wt-ICER transgenic zebrafish, referred to as 446A and 447A, respectively. Brightfield and fluorescence microscopy confirmed the successful establishment of the EGFP cell line, with robust EGFP expression observed in 446A cells (Fig. 4A). PCR analysis verified the integration of the HA-ICER transgene into the genome of the 447A cell line (Fig. 4B), and Western blotting further confirmed the expression of HA-tagged ICER protein in 447A cells (Fig. 4C). Notably, the presence of the N-terminal HA-tag likely prevents recognition by the ubiquitin-proteasome system, thereby stabilizing ICER protein and enabling its detection in this context. Immunocytochemical staining of 447A cells revealed nuclear and cytosolic localization of the HA-ICER protein, consistent with its function as a transcription factor and in conjunction with known delocalization of mono-ubiquitinated ICER from the nucleus (Healey et al., 2013). (Fig. 4D). To assess cell growth, we measured proliferation over five days. Both cell lines demonstrated comparable growth rates under standard culture conditions, though the HA- ICER cell line appears to divide faster, though not significantly (p > 0.05; Fig. 5).

**Figure 4.**
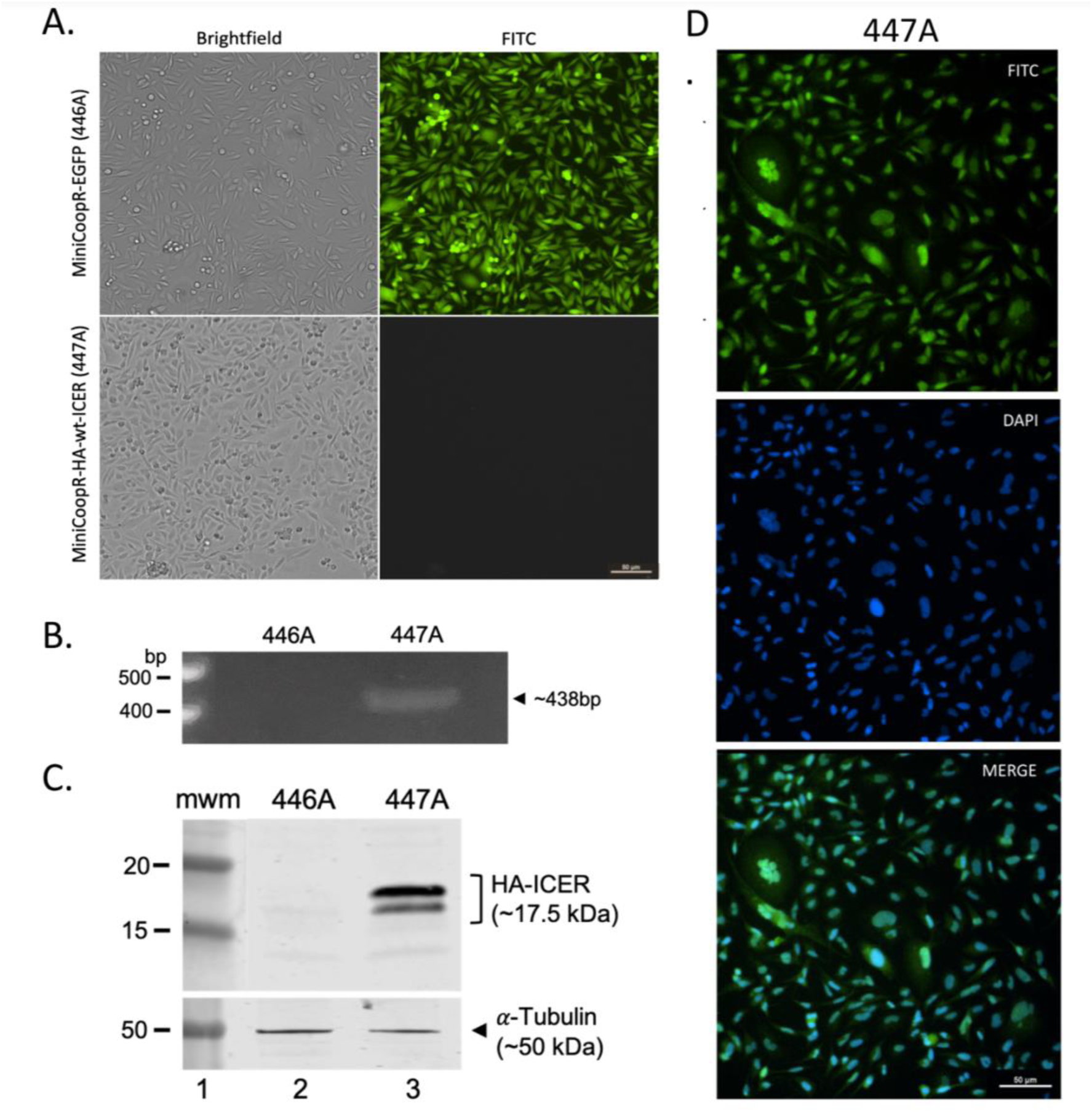
Characterization of established cell lines from fish melanomas. **A.** Images of brightfield and fluorescence microscopy of MiniCoopR-EGFP (446A) and MiniCoopR-HA-wt-ICER (447A) cell lines in culture. **B.** PCR analysis of 446A and 447A cell lines to test for HA-ICER transgene integration into the fish genome. **C.** Anti-HA Western blot analysis of 446A and 447A cell lines to test for transgenic HA-ICER protein expression. 446A was used as a control. **D.** Immunocytochemical determination of the subcellular localization of the transgenic HA-ICER protein in 447A cell line. 446A was used as a control, showing no signal (not shown). Commercially available anti-HA antibodies were used in the experiment shown in C and D, and anti-⍺Tubulin antibodies in C as loading control.

**Figure 5.**
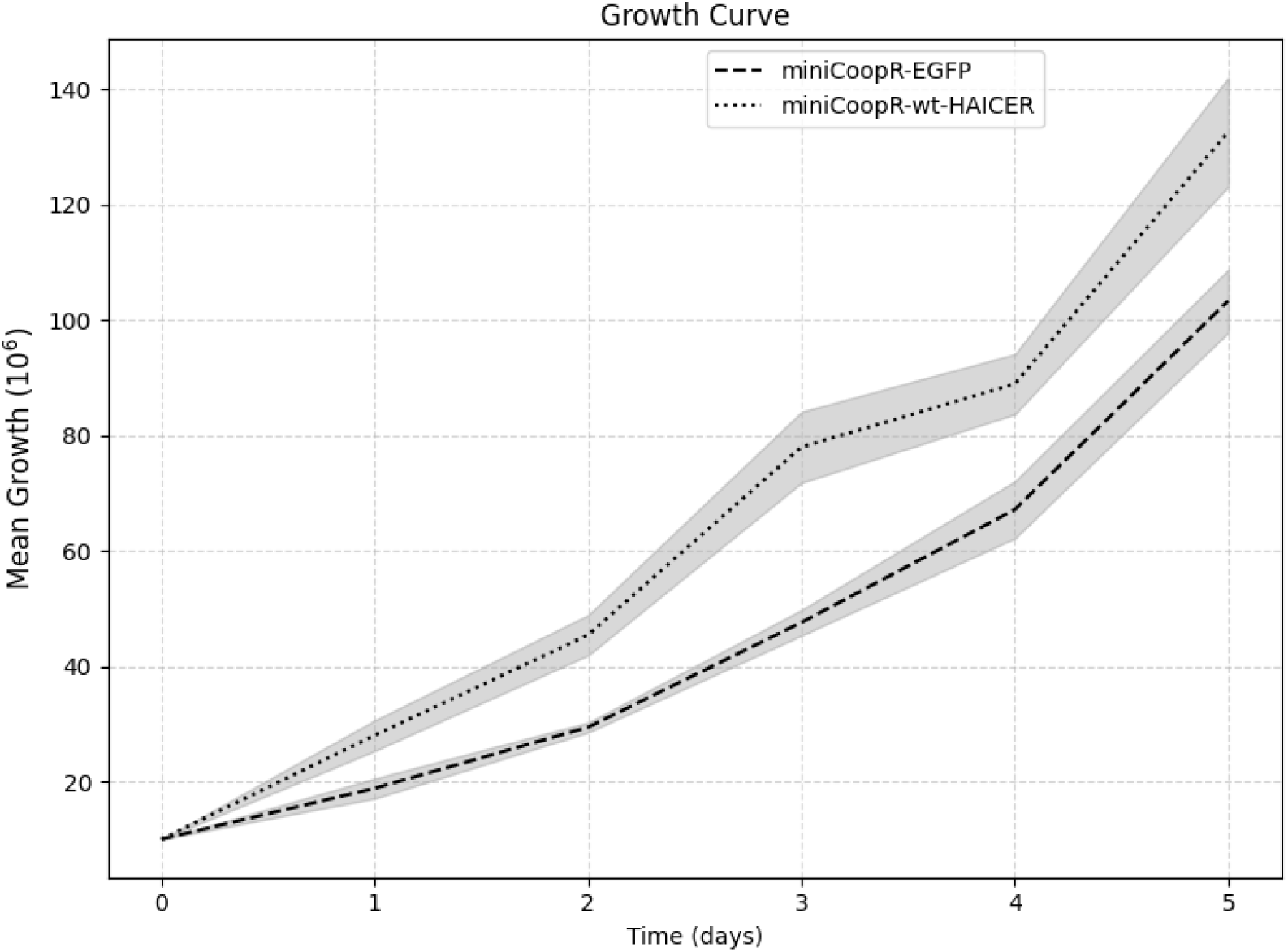
Cell growth curve. Cell number of in the indicated cell clones was monitored for five days. The values shown are the average of three independent determinations. Error bars (-/+) represent standard error. Students t-test was used to determine significance (p >0.05).

### Differential Gene Expression and Pathway Enrichment Analysis Reveals Tumorigenic Profile of wtICER

Using the previously validated cell lines derived from the zebrafish tumors, we sought to examine the transcriptomic profiles of the tumors. Three independent cultures per genotype were grown until about 1x 10^6^ cells per dish and total RNA was isolated to understand not only differentially expressed coding genes (DEGs), but also other non-coding RNAs that may be present in the sample. We found 2284 significantly (p<0.05) differentially expressed genes between the EGFP and wtICER fish tumor cells. DEGs were then analyzed using ShinyGO for Gene Ontology, which suggests pathways associated with more aggressive tumors in the wtICER were differentially expressed, in agreement with previously observed survival and histological studies. Of note, we observed differential expression of genes involved in glycosaminoglycan keratan sulfate biosynthesis (Fig. 6A), which has been implicated in melanoma and other cancers to contribute to focal adhesion and enhanced invasiveness (Tachibana et al., 2022; Leiphrakpam et al., 2019). Specifically, related to this pathway, we observed down-regulation of genes such as *b4galt3*, *st3gal1*, and a notable 5-fold increase in *b3gnt7* expression, a combination that is consistent with current metastasis literature in many cancer types (Chen et al., 2023; Wang et al., 2024). Other pathways shown in Figures 6B and 7A, contain overlapping genes, that have considerable literature about their interactions and relation to many types of cancer as well as melanoma, including but not limited to *akt1* (Cho et al., 2015), *cdkn1a* (Kadrmas et al., 2012), *col4a4* (Shin et al.,2022), *fgf5* (Ghassemi et al., 2017), *fgfr2* (Gartside et al., 2009), *met* (Zhou et al., 2019), *notch3* (Howard et al., 2013), *pdgfra* (Sabbatino et al., 2014), *pdgfrb* (Kadrmas et al., 2020), as well as upregulation of the proto-*oncogene junb*, which has also been shown to increase tumor growth and metastasis in murine models, and has been associated with poor outcomes in clinical breast and ovarian cancer patients (Pérez-Benavente et al., 2022).

**Figure 6.**
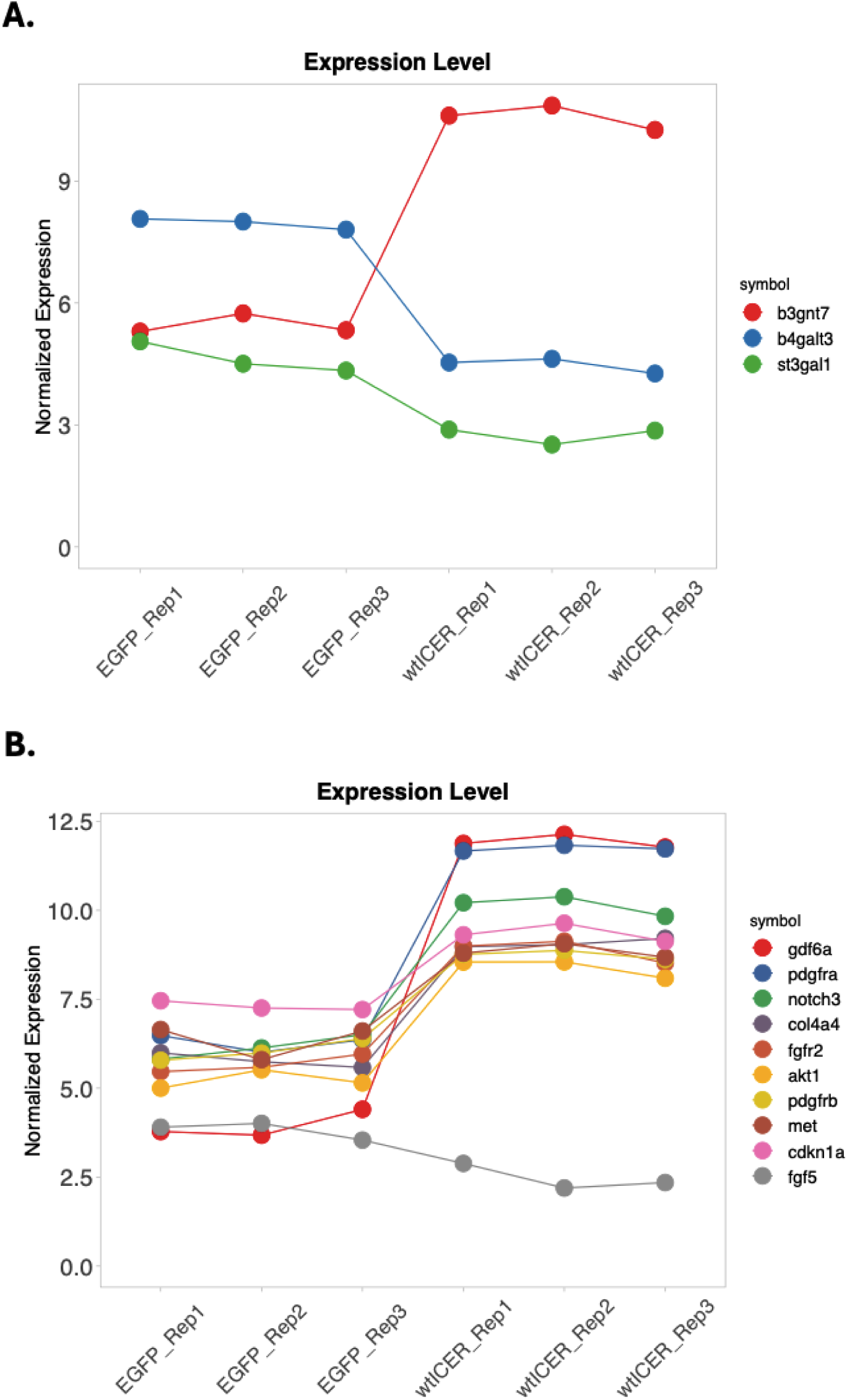
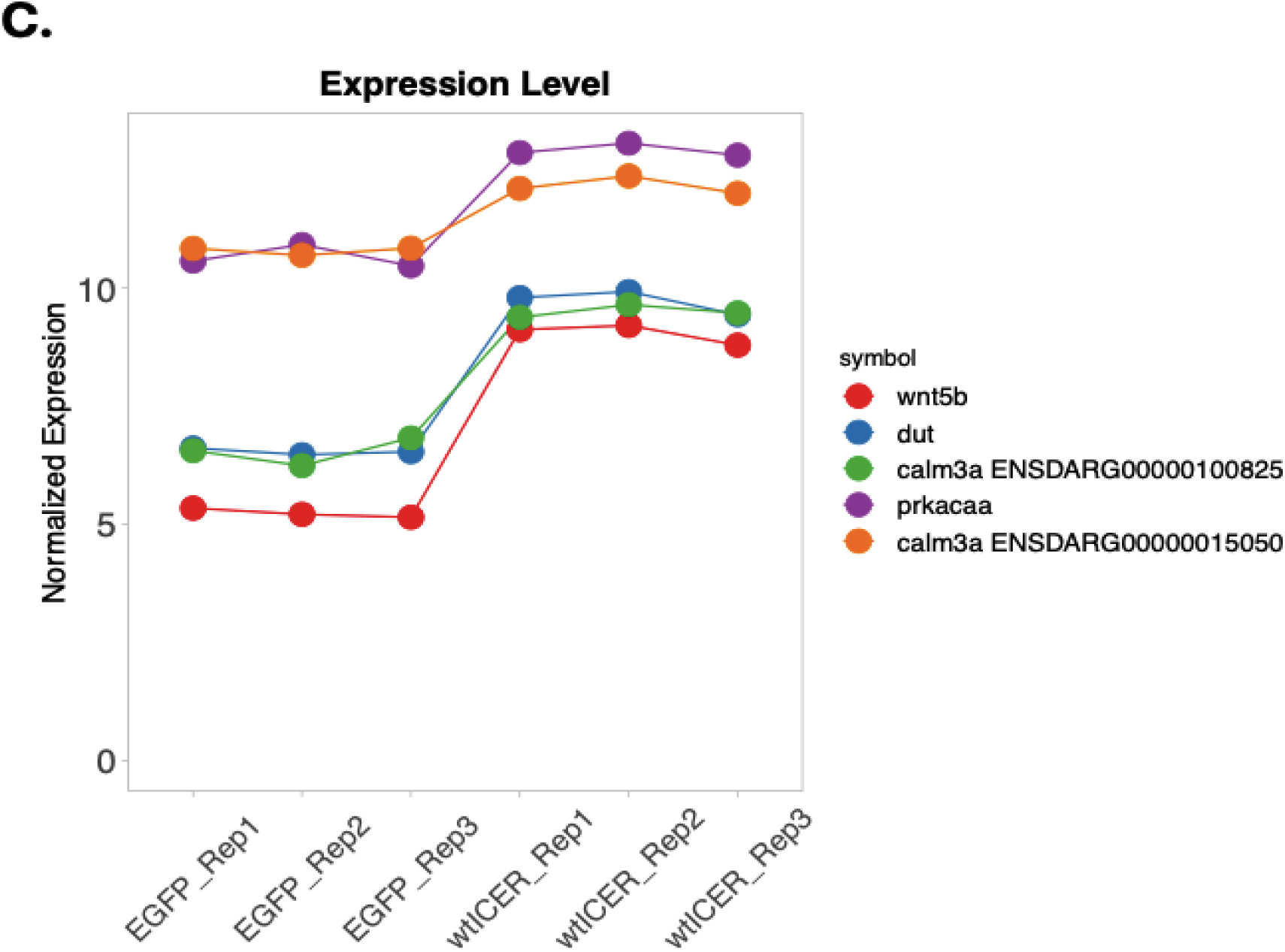
**Gene Expression comparison**. Expression signature of wtICER (n=3) and EGFP (n=3) of relevant genes involved in **A**. Glycosaminoglycan biosynthesis of keratan sulfate. **B.** Genes related to numerous tumor biology pathways, found to be differentially expression in this experiment. **C.** Genes found to be differentially expressed, based on DAVID query of possible ICER regulatory genes via Gene ontology, found to be involved in melanogenesis.

**Figure. 7.**
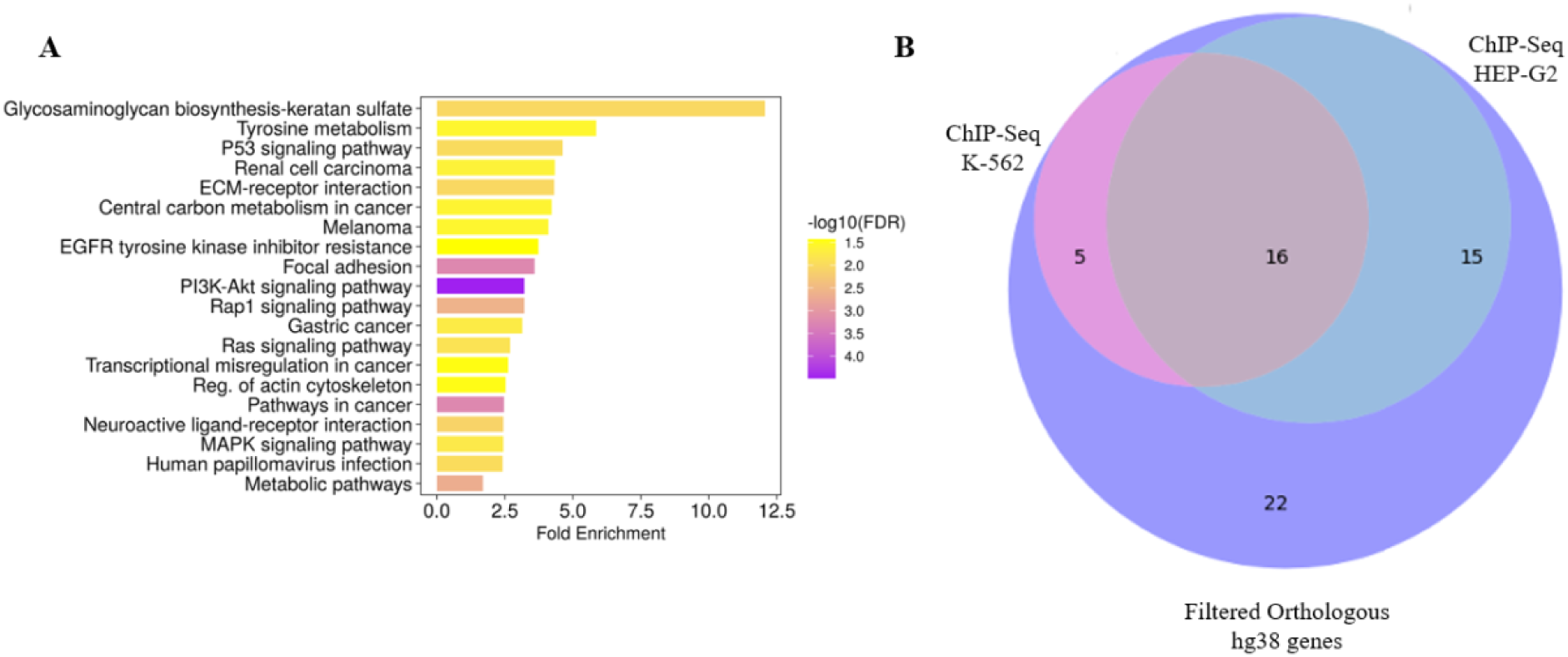
Pathway enrichment. **A.** Bar chart of ShinyGO output of all DEG from the zebrafish tumor cell lines for EGFP (n=3) and wtICER (n=3). We note that Glycosaminoglycan biosynthesis of keratan sulfate, while the highest Fold-enrichment, PI3K-Akt includes the lowest FDR, and includes many relevant genes involved in melanoma and other cancers including *akt1, cdkn1a, col4a4, fgf5, fgfr2, met, pdgfra, pdgfrb.* **B.** Venn diagram illustrating the overlap of genes identified through analyses of ICER’s transcriptional mechanisms. Total filtered list of 58 zebrafish and orthologous Human genes that were upregulated in wtICER, but downregulated in EGFP and phosphorylation mutant, and also contain CRE motifs in the promoter regions. 16 total genes overlap between 2 ChIP-Seq experiments from ENCODE (*crema, zcchc24, prkacaa, iqsec1b, rgs12b, sik2b, gli2a, gab1, ttyh3b, gyg1b, zbtb16b, actn4, junba, plxdc2, plpp3,* and *hmga2)*.

We also note an 8-fold increase in expression of the growth differentiation factor 6 *gdf6a*, which has been shown to be transcriptionally upregulated in melanoma and is involved in BMP signaling, promoting a neural crest signature and impacting differentiation (Venkatesan et al., 2018).

### Mechanistic investigation of ICER function suggests a link to auto-regulatory interaction with critical genes involved in melanogenesis

We then sought to elucidate the mechanisms contributing to the more aggressive melanoma phenotype observed in the wtICER transgenic zebrafish compared to the EGFP control and phosphorylation-deficient mutant cohorts. To start, we sorted our DEGs for significantly upregulated genes (p < 0.05) in wtICER compared to EGFP, consistent with the hypothesis that gene *upregulation* in this context could result from dysregulation of ICER as a transcriptional repressor. The promoter regions of these genes were analyzed for full CREs (5’ TGACGTCA 3’) and half-CREs (5’ TGACG 3’/5’ CGTCA 3’) using FIMO (Find Individual Motif Occurrences) via MEME-suite (version 5.5.7) with a threshold of p < 0.001 (Grant et al., 2011). This yielded a list of candidate genes that ICER may regulate through DNA binding at CRE sites.

We also created bulk RNA-seq libraries from solid neoplastic tissue for each respective cohort: EGFP (n=6), wtICER (n=6), and the phosphorylation deficient S35-41A-ICER (n=6). The expression patterns were expectedly heterogeneous, but helpful to directionally identify and filter for genes that were expressed at lower levels in the phosphorylation-deficient ICER tumors compared to wtICER. Drawing on prior studies in murine and human systems that established conserved ICER mechanisms and function, we reasoned that pertinent pathways and gene regulation would also exhibit conservation in this zebrafish model. To explore translational relevance, we identified the orthologous human gene IDs (hg38) for zebrafish genes using Ensembl biomart (Harrison et al., 2024) in our list and repeated the promoter analysis as described above. This cross-species analysis produced 58 genes with CRE motifs present in both zebrafish and human promoter sequences. These genes were upregulated in wtICER fish but not in the EGFP or phosphorylation-deficient ICER fish, suggesting a potential role for ICER dysregulation in their transcriptional activation.

To further support the functional relevance of these genes, we followed these analyses by analyzing publicly available Chromatin Immunoprecipitation and Sequencing (ChIP-seq) data from the ENCODE project. Specifically, we examined datasets targeting CREM in two distinct human cell lines: K-562, a chronic myeloid leukemia (CML) line derived from the pleural effusion of a 53-year-old woman with CML in blast crisis, and HEP-G2, a hepatocellular carcinoma line established from the tumor of a 15-year-old boy. Of the 58 candidate genes, 20 were found to overlap with CREM ChIP-seq peaks in the K-562 cell line, and 31 overlapped in the HEP-G2 cell line (Fig. 7B). We find 16 genes that overlap in both datasets (*crema, zcchc24, prkacaa, iqsec1b, rgs12b, sik2b, gli2a, gab1, ttyh3b, gyg1b, zbtb16b, actn4, junba, plxdc2, plpp3,* and *hmga2)*. Concordant ChIP-SEQ results coupled with our RNA-seq analysis provides evidence of ICER regulation, and antagonism of CREM/CREB, however, absent melanoma specific ChIP-seq data targeting CREM, it is unclear how cell type specific chromatin accessibility may impact ICER/CREM binding.

Gene Ontology (GO) analysis of the 16 candidate genes using DAVID (Sherman et al., 2022) revealed four genes *PRKACA, DCT, CALM3,* and *WNT5B* with key roles in the regulation of melanogenesis (Fig. 6C). These genes may be directly regulated by ICER via upstream binding to CRE motifs. Using UCSC genome browser, we manually confirmed that the promoter region for each gene contained CRE motifs (including half-CRE’s) based on the FIMO output, for both zebrafish GRCz11 and human hg38 genomes.

## Discussion

Although advancements in treatment such as immunotherapies and targeted therapies have improved patient outcomes in melanoma, there remains an urgent need for additional therapeutic strategies to address resistance and improve long-term survival. BRAF, and other MAPK pathway inhibitors, while initially highly effective at reducing tumor burden, are subject to drug resistance. One compensatory mechanism implicated in resistance to MAPK pathway inhibition, such as through BRAF and MEK inhibitors, is the activation of the cAMP signaling pathway. Intracellular cAMP levels can be elevated via G-protein-coupled receptor (GPCR) activation of adenylyl cyclase or through the inhibition of phosphodiesterase, which degrades cAMP. Increased cAMP activates Protein Kinase A (PKA), which phosphorylates various downstream targets, including members of the cAMP Response Element Binding (CREB) protein family. The Inducible cAMP Early Repressor (ICER), a potent transcriptional repressor and antagonist to CREB/CREM, plays a pivotal role in regulating tumorigenesis and has potential to mitigate resistance to BRAFi therapies. However, endogenous ICER’s therapeutic utility is inherently limited by its susceptibility to post-translational modification.

Phosphorylation at serine 41, mediated by MAPK, triggers polyubiquitination and proteasomal degradation, a process regulated by the N-end rule pathway and requiring recognition by the ubiquitin ligase UBR4, however N-terminal modification of ICER disrupts UBR4-mediated recognition, thereby preventing ubiquitination and degradation (Cirinelli et al., 2022).

Additionally, phosphorylation of ICER at serine 35 by the mitotic kinase CDK1 triggers monoubiquitination, resulting in its cytosolic relocalization. To circumvent both degradation and relocalization, we engineered an ICER mutant with serine-to-alanine substitutions at positions 35 and 41 (S35-41A-ICER) and included an N-terminal HA-tag to prevent N-terminal ubiquitin ligase recognition. This ubiquitination-resistant ICER mutant resists both mono- and polyubiquitination, retaining its nuclear localization and stability.

In this study, we investigated the impact of the ubiquitination-resistant ICER using a *Danio rerio* (zebrafish) melanoma model expressing *braf* V600E, *p53* loss-of-function, and *mitf* loss-of-function, rescued through the miniCoopR system. This model restores *mitf* expression only upon successful transgene integration, providing a robust system to study tumorigenesis in melanocytes. Histological analysis (H&E staining) revealed striking differences in tumor invasiveness among zebrafish expressing HA-tagged wtICER, S35-41A mutant ICER, and an EGFP control (Fig 3). The wtICER group exhibited the most aggressive tumor phenotype, while the S35-41A mutant ICER reduced tumor invasiveness. These findings suggest that the S35-41A ICER mutant sustains tumor-suppressive function, contrasting sharply with the highly invasive phenotype associated with wtICER. After generating the cohorts and a two-year observation period, we conducted a Kaplan-Meier survival analysis, which strongly suggests a survival benefit conferred by the S35-41A mutant ICER. Zebrafish expressing this mutant exhibited a demonstrable prolonged survival compared to the wtICER or EGFP control. Indeed, the difference in lifespan between all groups was statistically significant (p<0.05) (Fig. 2). While the S35-41A mutant ICER live longer than that wtICER and EGFP control, a larger sample size will be needed to increase statistical power to determine the impact on disease progression and survival in the S35-41A mutant phenotype. However, the accelerated mortality observed in the wtICER group correlated with aggressive tumor progression and clearly underscores the importance of ICER stability in tumor suppression.

Cell line experiments derived from tumors of these fish corroborated the histological and survival studies. wtICER-expressing cell lines exhibited faster, though not a significant, increase in proliferation compared to EGFP controls (p>0.05) (Fig 5). Interestingly, our attempts to establish stable cell lines via the same protocol for the S35-41A mutant were unsuccessful. We hypothesize the inability to grow the cells derived from this phosphorylation deficient fish’s neoplastic tissue is likely due to ICER’s constitutive impact on the cell cycle, driven by its intentionally impaired post-translational regulation and underscores the influence of ubiquitination-resistant ICER on cellular proliferation and warrants further investigation.

We also explore the gene expression profiles of the wtICER and EGFP tumor derived cell lines, which revealed several differentially expressed genes, with many pathologically relevant pathways to explore. We note many of interest, including the up-regulation of glycosaminoglycan keratan synthesis, as well as upregulated PI3K/AKT, Rap1 signaling, and upregulation of *gdf6*, which has been shown to be highly up-regulated in melanoma (Ceol et al ., 2011) however, the relationship, if any, between ICER and these pathways is not well understood. To begin to understand the mechanisms involved in promotion of a more aggressive melanoma in the zebrafish expressing wtICER, we focus here on an exploration of wtICER specific upregulated genes that have corresponding binding affinity for CREs, confirmed via ChIP-Seq data: *PRKACA* which encodes Protein Kinase A catalytic α-subunit and glioma-associated oncogene homolog 2 (*GLI2*). *PRKACA* was of interest due to its demonstrated implication in MAPK pathway inhibitors through its restorative phosphorylation and activation of CREB, and because the region surrounding the TSS contained the highest number of CRE motifs, which includes half-CREs (TGACG/CGTCA) amongst the melanogenic relevant gene set via DAVID. Resistance to MAPK pathway inhibitors has been shown to involve GPCR-mediated signaling, where GPCR activation induces adenylyl cyclase, leading to the production of cAMP and subsequent activation of PKA. Supporting this, the adenyl cyclase gene, *ADCY9,* and the PKA catalytic subunit alpha (*PRKACA*) were identified as key resistance effectors in melanoma, with *PRKACA* displaying the highest rescue score among serine/threonine kinases in a gain-of-function resistance study (Johannessen et al., 2013). Both *ADCY9* and *PRKACA* were demonstrated to confer resistance across a range of MAPK pathway inhibitors. These findings highlight a signaling network involving cAMP–PKA signaling as a driver of drug resistance in melanoma. Downregulation of *PRKACA* in a murine model reduced CREB Serine-133 phosphorylation and inhibited many pro-proliferative pathways (Wang et al., 2022). To our knowledge, it has not previously been suggested that ICER, as a transcriptional antagonist of CREB/CREM, is capable of inhibiting PKA-mediated phosphorylation of CREB, by downregulating *PRKACA,* the alpha catalytic subunit of PKA. The post-translational modifications of endogenous ICER, or the N-terminal HA-tagged ICER in this model, permit promoter CRE binding of CREB, which further stimulates PKA activity, and may partially explain the resurgence of phosphorylated CREB in drug-resistant melanoma. Our findings suggest that ICER may play a role in this signaling cascade by directly disrupting the cAMP– PKA–CREB feedback loop, by not only decreasing active CREB levels but also by mitigating the impact of the downstream targets of PKA. Here we posit a novel mechanism (Fig. 8) through which ICER could counteract resistance to MAPK inhibitors and potentially restore tumor sensitivity to therapy, though we also suspect similar mechanisms across a constellation of regulatory elements by ICER and other transcription factors contribute to this mechanism to fully inhibit melanoma progression. Further investigation is warranted to explore ICER’s regulatory role in this pathway and its therapeutic implications.

**Figure 8.**
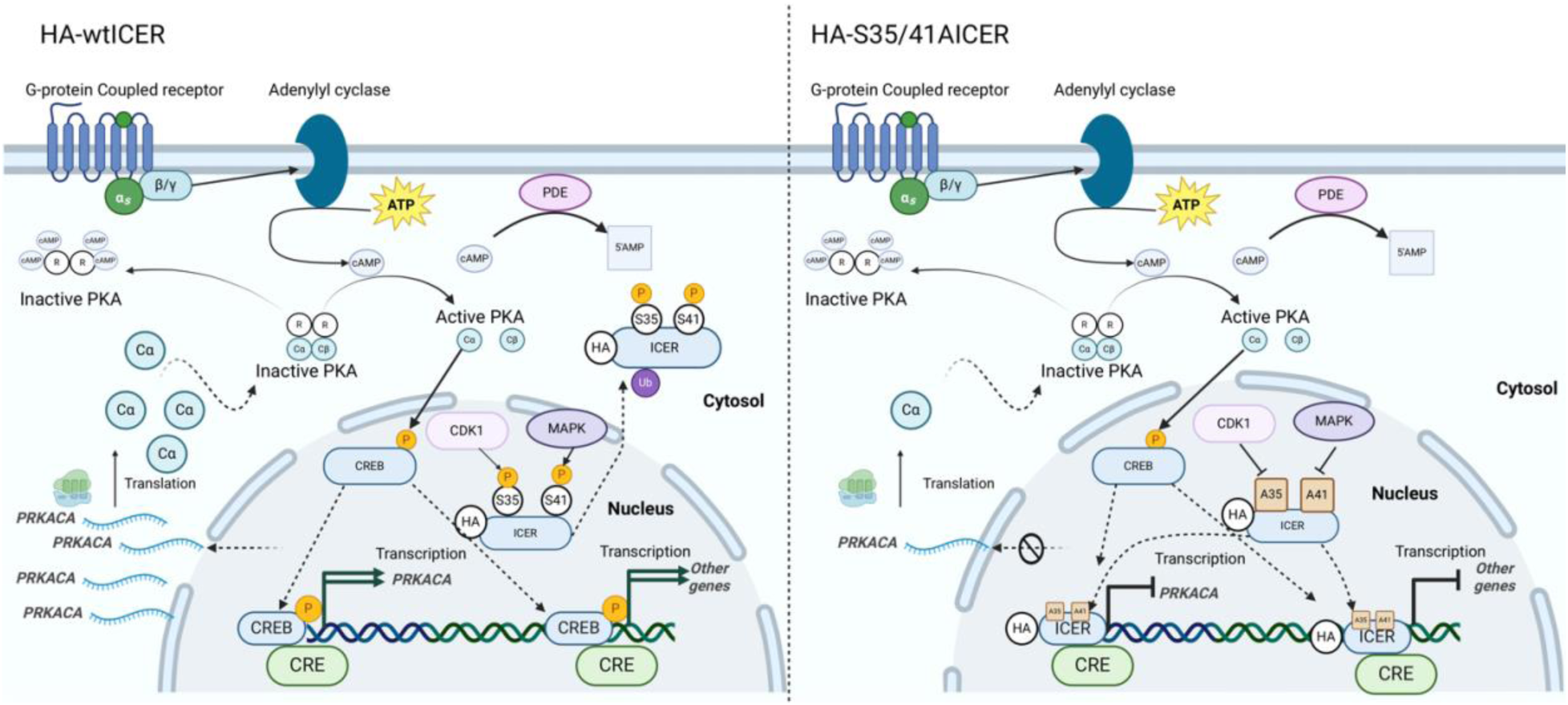
Proposed Model of PRKACA regulation via ICER Proposed model showing how wtICER (left) and the phosphorylation deficient S35-41-ICER (right) may impact gene expression, and specifically *PRKACA*, encoding the alpha catalytic subunit of PKA. Following adenylyl cyclase conversion of ATP to cAMP, 2 cAMP molecules bind to each PKA regulatory subunit, releasing the now active catalytic subunits, which phosphorylate CREB amongst other targets. Active CREB binds to CREs on the *PRKACA* promoter, upregulating gene expression of this subunit, which in turn, can more readily phosphorylate CREB in the presence of additional cAMP secondary messenger signal, forming a positive feedback loop. On the left, ICER is shown, in this case, to be phosphorylated on serine’s 35 and 41, and subsequently ubiquitinated and removed to the cytosol. ICER in the example shown is not poly-ubiquitinated or degraded due to the presence of an HA-tag which prevents N-terminal recognition by UBR4. On the right, we show the phosphorylation deficient ICER is now unable to be phosphorylated, and therefore not ubiquitinated. In this environment, ICER competitively binds to CRE’s on the promoter of many genes, including *PRKACA* and inhibits expression. In the case of *PRKACA* transcriptional repression by ICER, expression of the PKA catalytic subunit is decreased, decreasing the amount of phosphorylated and active CREB.

*GLI2*, an inducer of hedgehog (Hh) responsive genes, was of interest as another possible target of ICER regulation, because it also contains a high number of CRE motifs on the promoter region, and due to previous work highlighting the interest of this transcription factor in BRAF inhibitor (BRAFi) resistance. Notably, *in-vitro* melanoma cell lines with acquired BRAFi resistance show increased expression of *GLI2* and that knockdown of *GLI2*, restores sensitivity to vemurafenib (Faião-Flores et al., 2017). In other experiments, reduction in *GLI2* expression, inhibited both basal and Transforming Growth Factor-β (TGF-β) induced cell migration via matrigel assay, suggesting that the upregulation of *GLI2* contributes to a more invasive phenotype (Vasileia-Ismini et al., 2010; Faião-Flores et al., 2017). *GLI2* upregulation in melanoma was also associated with mesenchymal transition, which promotes cell-cell adhesion and the other motile behavior and is typically associated with poor prognosis (Javelaud et al., 2012). Interestingly, *gli2a* induced upregulated genes, such as *gli1* and *ccnd1*, were not identified as differentially expressed in our RNA-seq data. We hypothesize that while CREB/CREM can bind to motifs on the *gli2a* promoter and enhance its transcription (and ICER inhibits this process), the cells remain in a Hedgehog (Hh)-inactive state. In this state, PKA phosphorylates gli2a, leading to proteasomal degradation of the protein (Li et al., 2017). This degradation leaves only the transcriptional repressor domain active, which, in the absence of Hh signaling, causes no significant change in transcriptional activity.

Collectively, our results indicate that the degradation of wtICER disrupts its tumor- suppressive function, accelerating melanoma progression in this zebrafish model. In contrast, the ubiquitin-resistant S35-41A ICER mutant retains tumor-suppressive activity, evidenced by decreased tumor invasiveness and improved survival. We show that the autochthonous wtICER expression in melanocytes in a *braf^V600E^*, *p53*(lof) context leads to upregulation of metastatic- promoting genes involved in *PI3K/AKT/RAP1* signaling and glycosaminoglycan keratan sulfate synthesis, as well as progression towards a less differentiated melanocyte phenotype.

Additionally, we observe increased expression of key regulators of cell signaling and division, including *pdgfra*, *met*, and *notch3*. However, a direct mechanistic link between these pathways and ICER remains unclear, raising the possibility that resistant cells are being selectively enriched, or that additional, yet-uncharacterized mechanisms are at play. We offer some insight into the consequence of a ubiquitin deficient ICER, and what genes/pathways may be selected for in response to this environment, but more work will be needed to elucidate *how* a ubiquitin- deficient ICER can inhibit tumorigenesis.

## Methods

We want to thank Drs. Leonard Zon, Charles Kaufmans, Maurizio Fazio, Isabelle Roszko for helping establish our zebrafish facility, help with embryo injection techniques and general pathological analysis. Drs. Julien Ablain and Richard White for helping establish the fish melanoma cell lines. Dr. Laying Wu for tissue processing and microscopy.

## Funding

This work was supported by the National Institutes of Health [grant number 5SC1GM125583] Zebrafish were maintained in accordance with the guidelines set forth by the Institutional Animal Care and Use Committee (IACUC) to ensure ethical and humane treatment of vertebrate animals in research. All experiments involving zebrafish were carefully designed to minimize pain and distress, utilizing appropriate anesthetics and analgesics when necessary. Housing and care comply with the Guide for the Care and Use of Laboratory Animals, including maintaining zebrafish in well-regulated aquatic facilities with optimal water quality and enrichment. Protocols undergo rigorous review and approval by the IACUC, ensuring that the principles of the 3Rs—Replacement, Reduction, and Refinement—were upheld throughout our research endeavors.

## Notes

### Competing Interest Statement

The authors have declared no competing interest.

